# Toward Health Management of Major Labour Force Generation by Using Infection Control Countermeasures for Haematobium Schistosomiasis – assumed to be related to occupational risk-in the Republic of Malawi

**DOI:** 10.1101/535542

**Authors:** Nobuyuki Mishima, Samuel K. Jemu, Tomoaki Kuroda, Koichiro Tabuchi, Andrew W. Darcy, Takaki Shimono, Pheophet Lamaningao, Mari Miyake, Seiji Kanda, Susan Ng’ambi, Yoshihiro Komai, Hirofumi Maeba, Hiroyuki Amano, Toshimasa Nishiyama

## Abstract

**Background:** In Malawi, haematobium schistosomiasis is highly endemic. According to previous studies, countermeasures have been conducted mainly in school-aged children. In this study, we focused on the age groups, which are assumed to be major labour force generation. Haematobium schistosomiasis is supposed to be related to occupational activities in schistosome endemic countries.

**Methods:** We chronologically followed the transition of schistosome egg positive prevalence before and after mass drug administration of praziquantel (MDA) by using a urine filtering examination. We also analyzed the effectiveness of urine reagent strips from the cost perspective.

**Findings:** The egg positive prevalence was 34.3% (95%CI: 28.5-40.5) just before MDA in June 2010 and the highest prevalence was in the age of twenties. The egg positive prevalence reduced to 12.7% (95%CI: 9.2-17.3, p<0.01) eight weeks after the first MDA and the prevalence reduced to 6.9% (95%CI: 4.6-10.0, p<0.01) after the second MDA in August 2011. The egg positive prevalence after MDA in 2013 was reduced from 3.8% (95%CI: 2.1-6.9) to 0.9% (95%CI: 0.3-3.4) and p value was 0.050. Using urine reagent strips after MDA, the positive predictive value decreased, but the negative predictive value remained high. The cost of one urine reagent strip and one tablet of praziquantel were US$0.06 and US$0.125 in 2013 in Malawi. If the egg positive prevalence is 40%, screening subjects for MDA using urine reagent strips, the cost reduction can be estimated to be about 24% -showing an overall cost reduction.

**Conclusion:** The combination of MDA and urine reagent strips could be both a practical and cost-effective countermeasure for haematobium schistosomiasis. It is key to recognize that haematobium schistosomiasis could be considered a disease that is assumed to have some concern with occupational risk in tropical agricultural countries such as Malawi. From this point of view, it is very important to protect the health of workers; the sound labour force generation is vital for economic growth and development in these countries.

**Author summary:** Schistosomiasis is widely endemic in the tropical and subtropical countries including Malawi, and it is related that more than 300 million people suffer from associated severe morbidity. The pathway of transmission is mainly contacting infested fresh water and it is inevitable to contact fresh water through their daily activities in Malawi. Then, they are routinely exposed to the risk of schistosome infection. Previously the main targets of schistosome control were school-aged children, but our research showed main population of schistosome infection was twenties that was presumed to be major labour force. Agriculture is the dominant industry in Malawi and it can be related to be at risk of schistosome infection during agricultural work. Schistosomiasis is presumed to have occupation-related risks, we consider that schistosome control will be a valuable step-up to economic development and make a social contribution in Malawi and many low-income tropical countries.

**Funding:** The Ministry of Education, Culture, Sports, Science and Technology of Japan’s scientific research grant (<u>JSPS KAKENHI Grant Number JP23406025</u>). The funders had no role in study design, data collection and analysis, decision to publish, or preparation of the manuscript.

## Introduction

Schistosomiasis, a trematode infectious disease, is widely distributed around the tropics and subtropics. This infectious disease is one of the world’s three major parasitic infections. It is endemic in 74 developing countries; and approximately 800 million people are at risk of schistosome infection. More than 300 million people suffer from associated severe morbidity [1]. Chronic and repeat infection of schistosomiasis could result in irreversible damage to body organs and other diseases; for example, *Schistosoma haematobium* infection may lead to bladder cancer and cervical cancer [2, 3]. In the schistosome endemic regions, the most prevalent form of the disease is chronic schistosomiasis, resulting from repeated exposure to infectious cercariae [4]. Schistosomiasis mortality rates rises substantially as age increases [5]. Therefore, it is important for healthcare systems to consider not only children but also young adults –assumed to be a major component of labour generation-as the subjects of schistosomiasis control as related to occupational risk.

Schistosomiasis is recognized as one of the neglected tropical diseases (NTDs) at present. Global coverage rate of preventive chemotherapy against schistosomiasis is still low at 8.3%, while the rate against onchocerciasis is 59.8% [1]. In sub-Saharan Africa, approximately 280,000 annual deaths have been attributed to schistosomiasis [6]. Countermeasures have been globally to fight malaria, tuberculosis, and HIV infection; however, we consider provisions against schistosomiasis are an important next step for the sustainable growth and development in countries affected by schistosomiasis because of the burden the disease places on those living in the affected region. This disease does not only cause immediate morbidity in children, but it also has long-term health effects on the children’s development into adulthood. Although the earlier studies targeted school-aged children, but the disease may become a matters of concern for the general public if the overall labour force suffers from schistosomiasis.

Malawi is one of the poorest countries in the world and was ranked as the ninth poorest country in terms of GDP per capita in 2009 that stood at US$290 [7]. And it is still US$300.79 in 2016 on the list of World Bank. Malawi is an endemic country of schistosomiasis and schistosomiasis is associated with populations living in poverty in sub-Saharan Africa, including Malawi. In order to ameliorate poverty, it is important to improve the working environment; and managing the health of the working population is very important to establish a stable and sustainable economic environment. From the viewpoint of labour healthcare management, controlling schistosomiasis could be one of the most effective countermeasures for those countries affected by schistosomiasis. There are few researches that have taken measures against schistosomiasis from the viewpoint of occupational risk.

In Malawi, the main pathogens are *Schistosoma haematobium* and *S. mansoni*. Schistosomiasis is transmitted through contact with infected freshwater in which these intermediate hosts live. The two intermediate hosts distribute simultaneously. *Bulinus globosus*, the main intermediate host, is distributed all over the country-especially where there are sources of freshwater, such as sugarcane plantations, rice growing schemes, man-made dams, rivers, and ponds. While the genus *Biomphalaria* occurs in Lilongwe and the Linthipe Plain, Chapanaga area in Chinkhwawa district; some parts of Ntchisi, Salima, Karonga, Namwera in Mangochi, and Blantyre. It is reported that schistosomiasis is more rampant in poor rural communities especially places where fishing and agricultural activities are dominant [8]. The pathway of schistosome transmission notably affects farmers, fishermen, irrigation workers, and those whose daily activities involve contact with infested freshwater. Contact with freshwater is the inextricable part of the daily activities of many inhabitants in the area. Malawi is predominantly an agricultural country and agriculture accounts for about 35% of GDP. Moreover agricultural activities provide more than 80% of the employment in this country [7]; therefore, the vast majority of the population is routinely exposed to the possibility of schistosome infection while working. As a result of this situation, we need to recognize that there could be the occupational risk in suffering from schistosomiasis. The prevalence of the disease in the country is estimated between 40% and 50%; school-aged children are a highly infected group and are intensely affected [9]. It was previously reported that although all sections of the population in the endemic areas can be infected with schistosomiasis, the most vulnerable groups are pre-school (under 5 years old) and school-aged children, adolescent girls, and women of childbearing age [10, 11]. Haematobium schistosomiasis is likely to impact child growth and possibly can cause anemia in all age groups; this would call for the inclusion of the entire populations into future control programs [12].

In this study, which looks at schistosomiasis that could have relation to occupational risk, we targeted residents of all generations, including major segments of the labour force to check the current status of residents in our surveyed areas using mass drug administration of praziquantel (MDA) and urinalysis. Protecting the health of the labour force can be expected to contribute to the economic growth and development of Malawi. Findings from our study should also help other tropical and subtropical schistosome endemic countries.

## Methods

### Study area and population

A survey was conducted in twelve contiguous villages with similar socio-economic and cultural characteristics in Nkhotakota District located on the shores of Lake Malawi from June 2010 to August 2011. The total population surveyed was 1,810 people, more than 4 years old. Around 300 subjects were selected by random sampling for urine examinations. I threw a Japanese 10 - yen coin and chose the participants who got the head of a coin in order. The inhabitants of these villages were predominantly subsistence farmers of maize and rice and river/swamp fishermen. The communities had one primary school and functional boreholes. The people can access a healthcare center by walking a few hours. Schools provided health education for the prevention of schistosomiasis.

In 2012 a second survey was conducted. Four target areas in the Lilongwe District were selected where previously we had conducted mass drug administration using praziquantel for all the residents more than 4 years old in 2012 with the help of the Malawi Ministry of Health (Community Health Science Unit). These four target areas were Chisindo, Mtika, Mapiri, and Chisaka in Lilongwe. The total population was 1,393 people in these four areas, and around 300 subjects were selected by random sampling from among all age groups. The target areas were located near the capital Lilongwe, the basic infrastructure was being developed to some extent. Health education for schistosome infection had been provided at the schools. Information about schistosomiasis was also provided via broadcast media.

### Urine examination survey

A list of registered inhabitants (1,810 people) was prepared by a door-to-door survey in April 2010. In June 2010, 242 participants were identified; then 260 people in August 2010; 315 people in June 2011; and later 350 people in August 2011. We distributed instruction on urine examination written in Chichewa (Malawian domestic language) to all participants and then they received explanation on urine examination in Chichewa from the Malawi Ministry of Health staff. After finishing the questions and answers about the explanation more than one hour, only those who agreed to the examination were selected as the participants. We got the informed consents in writing by them. The participants provided their urine samples and with a urine test administered among randomly sampled subjects. In 2013, 264 people in June and later 211 people in August were among randomly selected subjects who participated in our urine examination survey. We examined all urine samples provided from all participants, regardless of age. The urinalysis was performed twice in a year; the first was done immediately before mass drug administration of praziquantel and the second analysis was performed 8 weeks after the administration [13]. The freshly passed mid-day urine samples were collected from 10 am to 2 pm, and were screened for microhaematuria and proteinuria using urine reagent strips (SD UroColor 11, Standard Diagnostics Inc., Korea). The urine reagent strips were used according to the manufacturer’s instructions and all strips were checked in about thirty seconds. The specimens were then processed for microscopic examination of schistosome eggs at the site of the collection within three hours.

Processing the specimen and egg detecting followed the syringe filtration technique. A urine subsample of 10mL was drawn into a plastic syringe from each well-mixed sample and strained through a nylon filter (12 μm pore size: Disease Control Textiles, Vestergaard Frandsen Group, Denmark). The filter was then examined under the microscope with magnification of x100, and *S. haematobium* eggs were detected.

### Mass drug administration

After the urine examinations were completed, we provided the information of praziquantel (E. Merck KG) written in Chichewa to all villagers. They received explanation on the drug in Chichewa from the Malawi Ministry of Health staff. Then we got the informed consent for administering praziquantel in writing from the participants. In the case of those participants who were under 20 years of age, we also obtained confirmation from their guardians. Concurrently, we checked, in advance, the drug allergy history of all the participants and no one had a praziquantel allergy. Regarding the safety of praziquantel administration, we consider those under the age of 4 years as an ineligible population. Women within their first trimester of pregnancy and those who had a history of epilepsy or other signs of potential neurological disease were also excluded from mass drug administration. Praziquantel 40mg/kg was administered to eligible participants (838 people in 2010 and 1,027 in 2011, 728 in 2013), who agreed by Directory Observed Treatment, Short Course (DOTS) protocol by using weight scales. After praziquantel administration by DOTS, we confirmed whether there were no adverse reactions to each participant for at least 30 minutes under the supervision of physicians. In addition, the Malawi Ministry of Health staff lectured on prevention of schistosomiasis to all participants using printed materials written in Chichewa.

### Ethical consideration

Clearance to undertake the follow up study was obtained from the District Health Office of the Ministry of Health (MOH) in Malawi and Kansai Medical University Ethical Committee in Japan. In the areas used in the study, permission to proceed with the study was obtained from the District Health Officer (DHO). The approved number of Ethics Review Committee (KAN-I-RIN) is 0758.

A signed consent form was obtained from both the new and old participants in the study. The consent form contained the following information: general introduction of the study, usefulness of the study, and purpose of the study. The participants were allowed to withdraw at any point during the project whenever they deemed appropriate.

### Data analysis

The quantitative data were analysed using SPSS (version 20.0.0). First, SPSS was used for cross tabulation and running of frequencies. Secondly, the chi-square test was used to establish the relationship between the categorical variables. A p value less than 0.05 was considered statistically significant. Graphs were drawn using Apple Inc. Numbers (Ver.4.3.1).

## Results

Table 1 shows the summary of urine filtering examination. The study focused on a random sample of people who participated in our survey: 242 people (age: 22.69±35.56 years) in June 2010; 260 people (age: 22.81 ±35.47 years) in August 2010; 315 people (age: 23.55 ±38.00 years) in June 2011; and 350 people (age: 23.56±37.57 years) in August 2011. Research showed the egg positive prevalence for males was 42.6% (95%CI: 33.6-52.1) and for female participants was 27.6% (95%CI: 20.7-35.8). Fig 1 shows the egg positive prevalence among all participants in 2010-2011. The schistosome-egg positive prevalence among the participants was 34.3% (95%CI: 28.5-40.5) just before mass drug administration (MDA); and this result was almost the same as the egg positive prevalence in Malawi that was previously reported. By age group, the highest egg positive prevalence was detected in participants in their 20s and it was 47.5% (95%CI: 32.8-62.6) before the first MDA, among this group the egg positive prevalence of males in their 20s was 53.3% (95%CI: 29.9-75.4). The first MDA was conducted for 838 participants (46.3%) just after the urine examinations. No one who took praziquantel showed any obvious side effects. The schistosome egg positive prevalence 8 weeks after MDA was 12.7% (95%CI: 9.2-17.3), and it was significantly lower than that before the mass treatment in June (p<0.01). Egg positive prevalence decreased in all age groups. One year after the first MDA, egg positive prevalence was 14.6% (95%CI: 11.1-19.0) in June 2011 and MDA supposedly kept the prevalence low. However, the prevalence increased in the under-15 age group shown in Figs 2-a and b, those are the process of egg positive prevalence by age groups in 2010-2011. Among those in the under-15 age group, the gradient of the increasing line of the under-5 group was steepest. The egg positive prevalence of those in the under-5 age group increased from 0% to 26.7% ten months after checking urinalysis in August 2010. On the other hand, the egg positive prevalence for participants between 15 and 19 years remained the same and the prevalence of those in the older than 20 age group decreased. The second MDA for 1,027 participants (56.7%) was conducted just after the urine examination in June 2011, and 8 weeks later the egg positive prevalence was 6.9% (95%CI: 4.6-10.0). The egg positive prevalence significantly decreased after the second MDA (p<0.01) in August 2011 and the total reduction rate completed sequential MDA was 79.9%. The egg positive prevalence was significantly reduced among those in the age groups of 10-14 year-olds and 15-19 year-olds (p<0.05). After the second MDA, none of the people who took praziquantel had obvious side effects.

**Fig 1.**
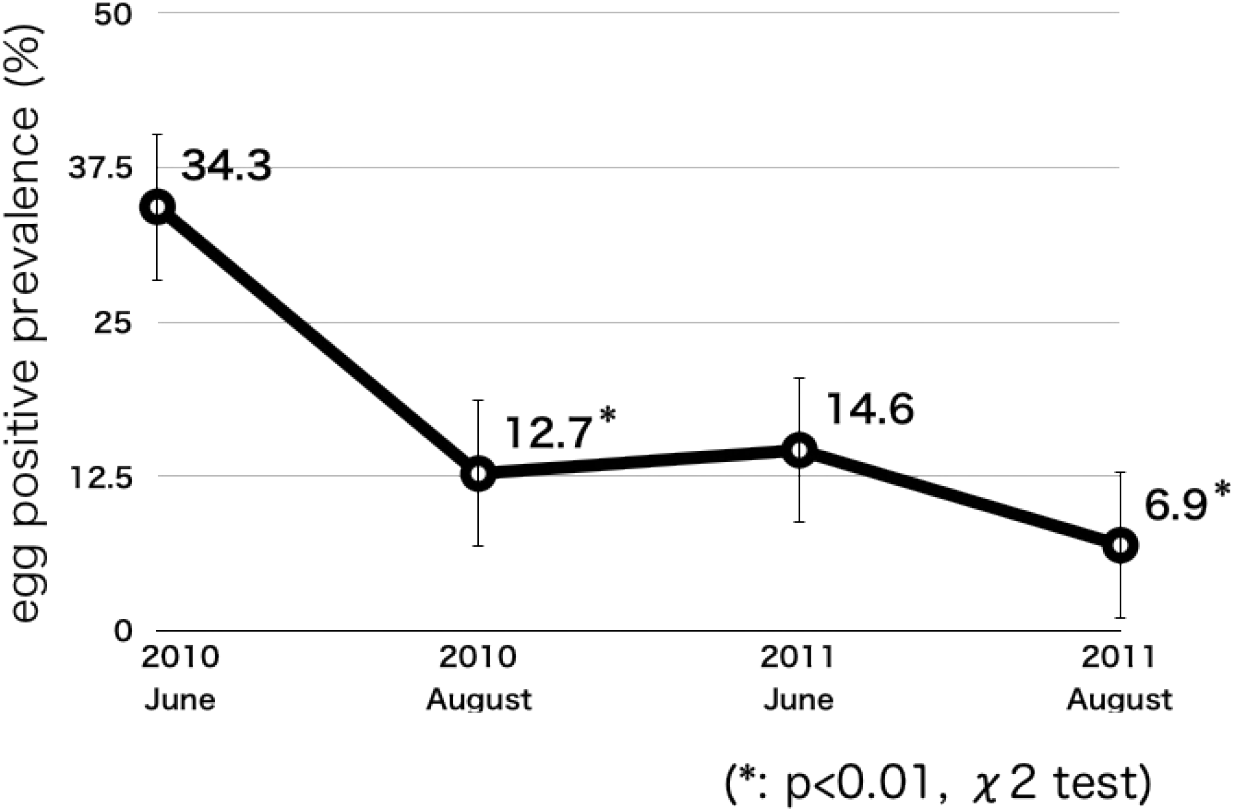
Process of egg positive prevalence among all participants 2010-2011

**Fig 2-a.**
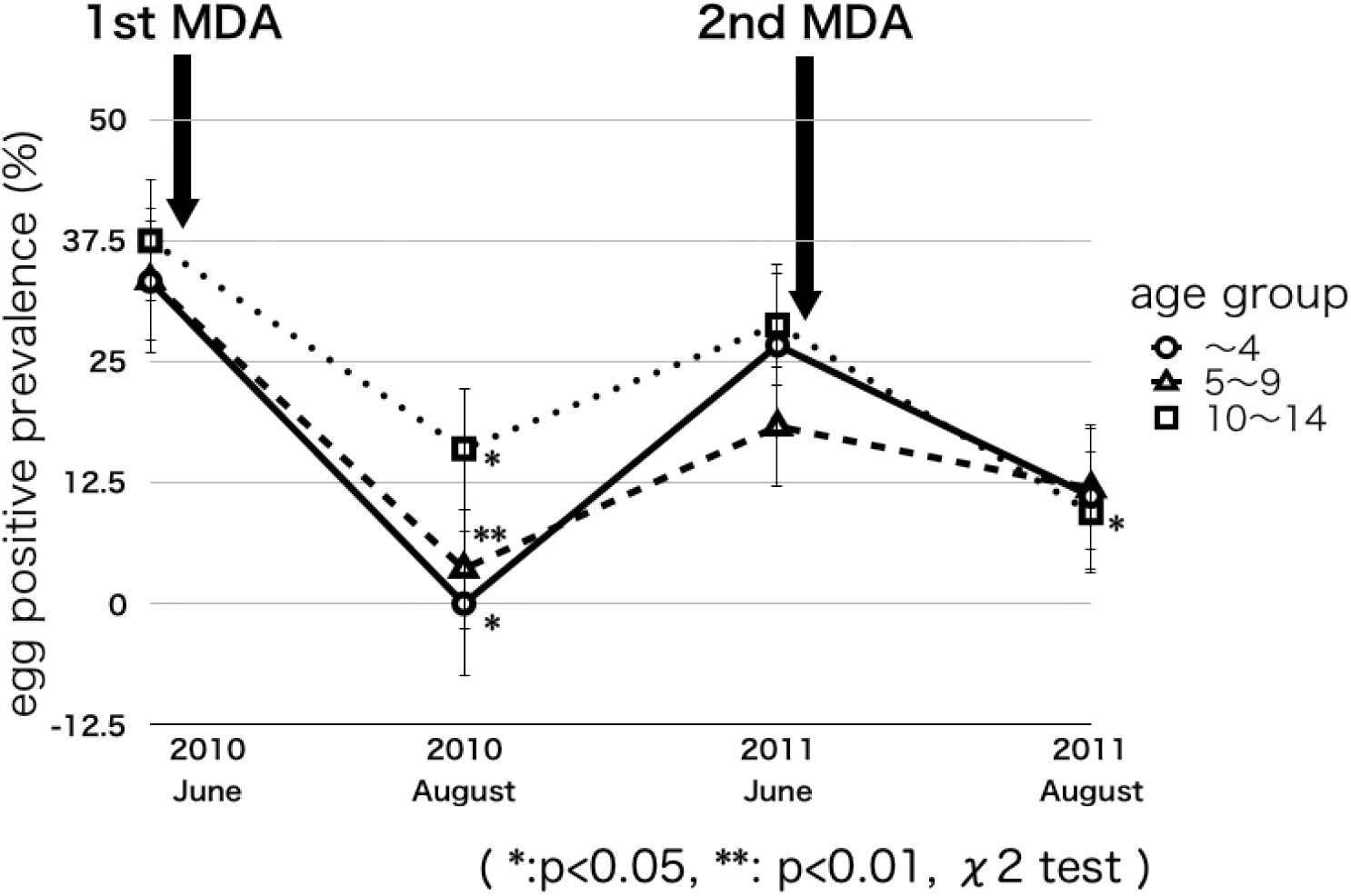
Process of egg positive prevalence by age group 2010-2011

**Fig 2-b.**
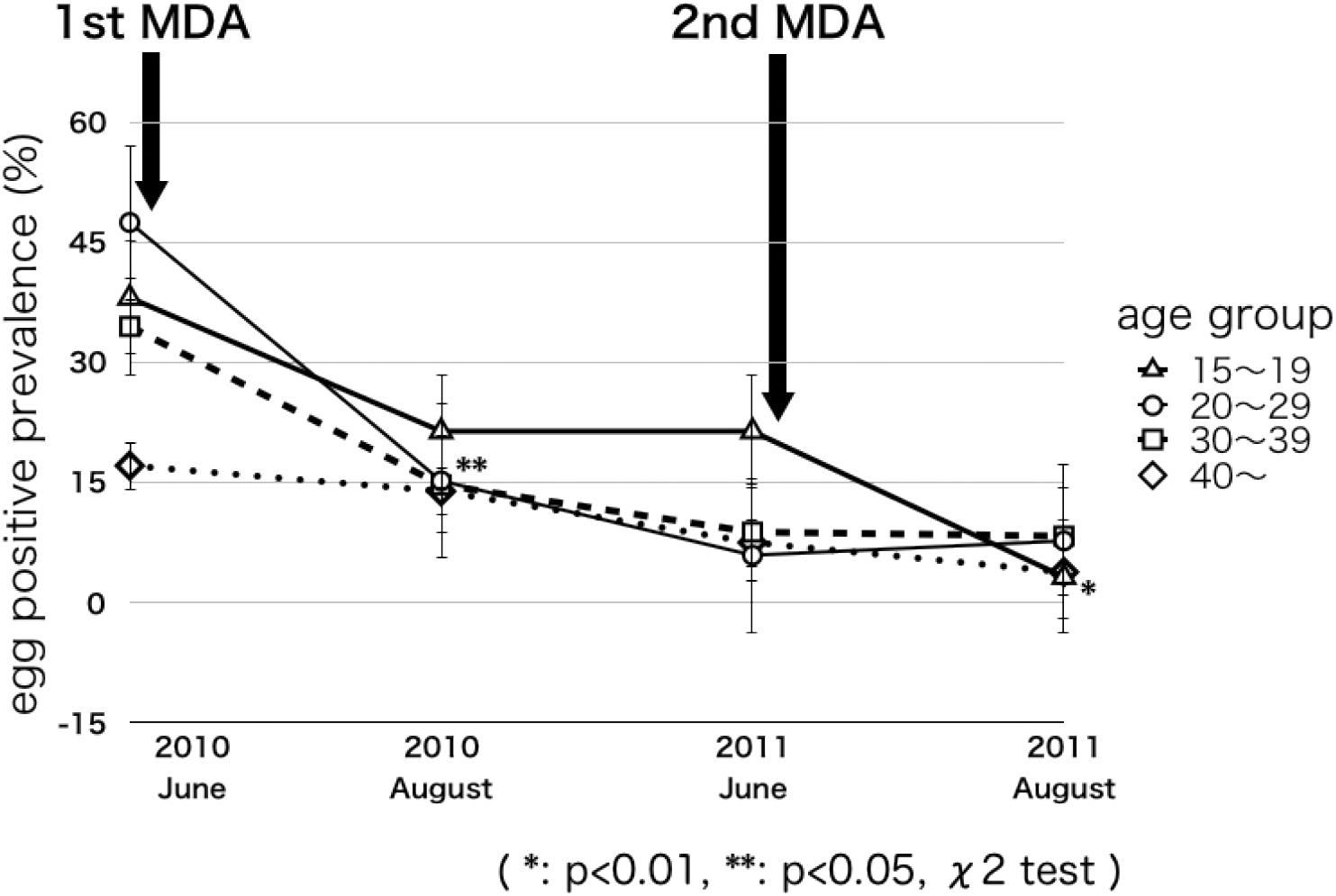
Process of egg positive prevalence by age group 2010-2011

**Table 1.**
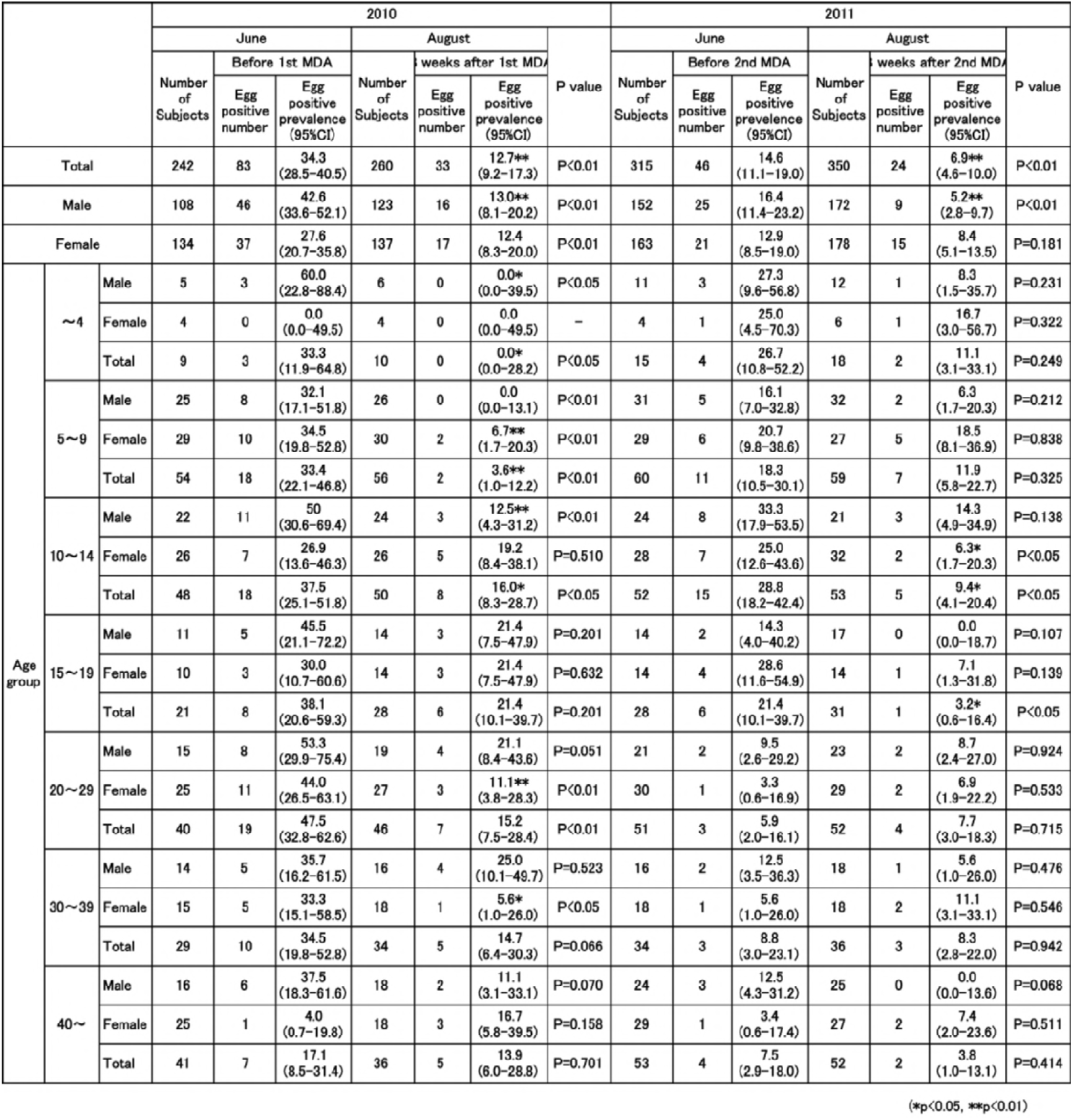
Results of urine filtering examination 2010-2011

Fig 3 shows analysis for the results of occult blood by the urine reagent strips in 2010-2011. After MDA, the positive predictive value decreased but the negative predictive value remained high, more than 96%. After the second MDA, the negative predictive value was 99.2% (95%CI: 98.1-100). The sensitivity was 91.7% (95%CI: 80.6-100) and the specificity was 87.7% (95%CI: 83.9-91.6).

**Fig 3.**
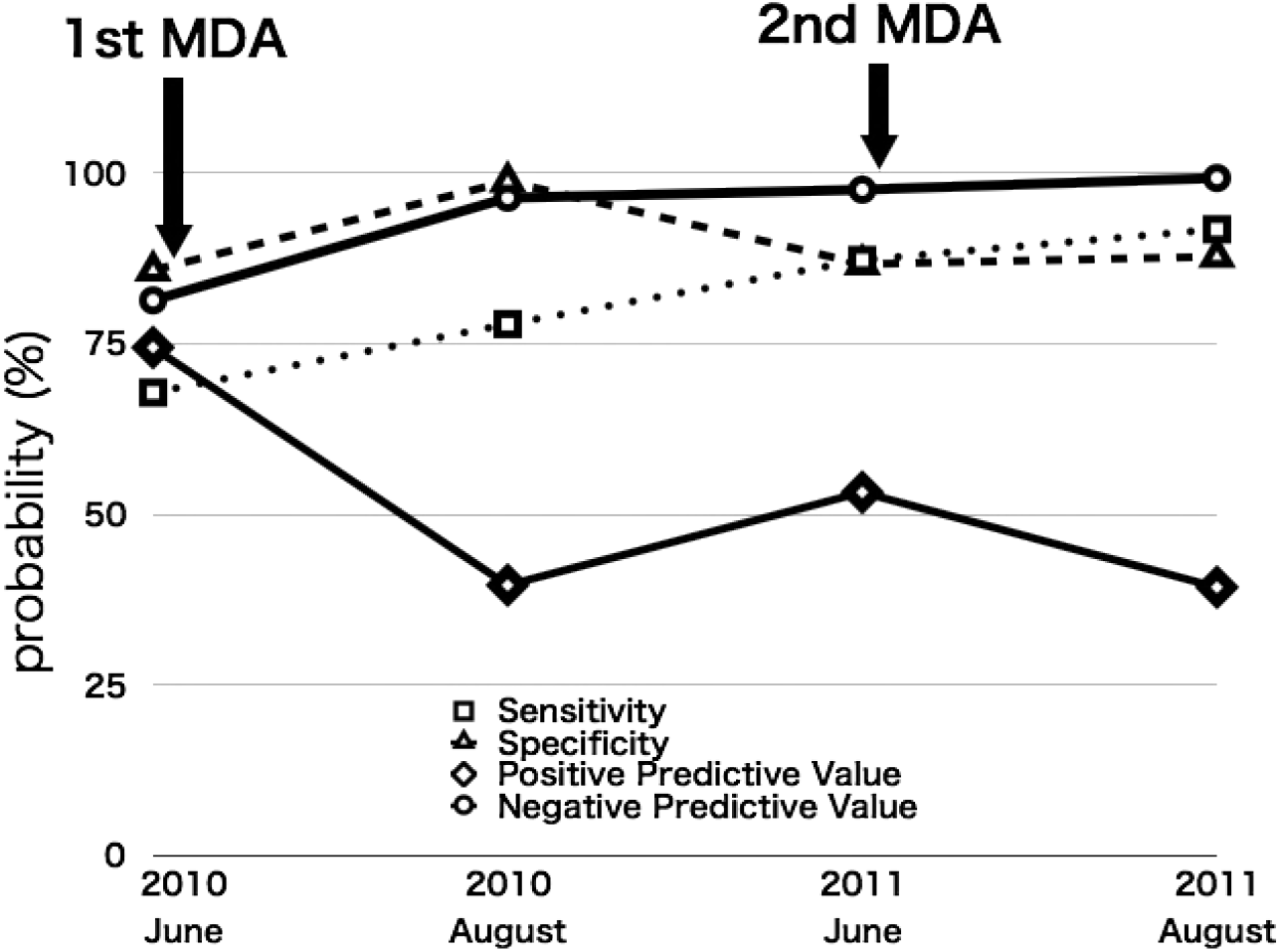
Analysis for the results of occult blood by urine strip tests

The surveyed areas in 2013 were the same areas where the MDA of praziquantel was previously conducted in 2012. Table 2 shows that overall MDA coverage rate in 2012 had ranged from 50.25% to 85.8%. Table 3 shows the result for the urine filtering examination in 2013. Total participants were 264 people (age: 21.78±38.42 years) in June and 211 (age: 19.99±36.66 years) in August; and they were selected by random sampling among all the residents. The egg positive prevalence in 264 participants was 3.8% (95%CI: 2.1-6.9), it was relatively lower than the average prevalence of the country and the highest was 8.8% (95%CI: 3.0-23.1) in those under five (Table 3). Eight weeks after MDA, egg positive prevalence decreased to 0.9% (95%CI: 0.3-3.4, p=0.050). The reduction rate was 76.3%. Schistosome eggs were not detected at all in most age groups, excluding the 15-19 age group. Those in the 15-19 age group had an egg positive prevalence was 11.1% (95%CI: 3.3-33.1) 8 weeks after MDA shown in Figs 4a and b, those are the process of egg positive prevalence by age groups in 2013. The youngest subject among the egg positive group was a 2-year-old boy. Praziquantel 40mg/kg was administered to 728 participants (52.3%) more than 4-years-old by DOTS immediately after urine examination. We confirmed that no one experienced any side effects from the praziquantel administration.

**Fig 4-a.**
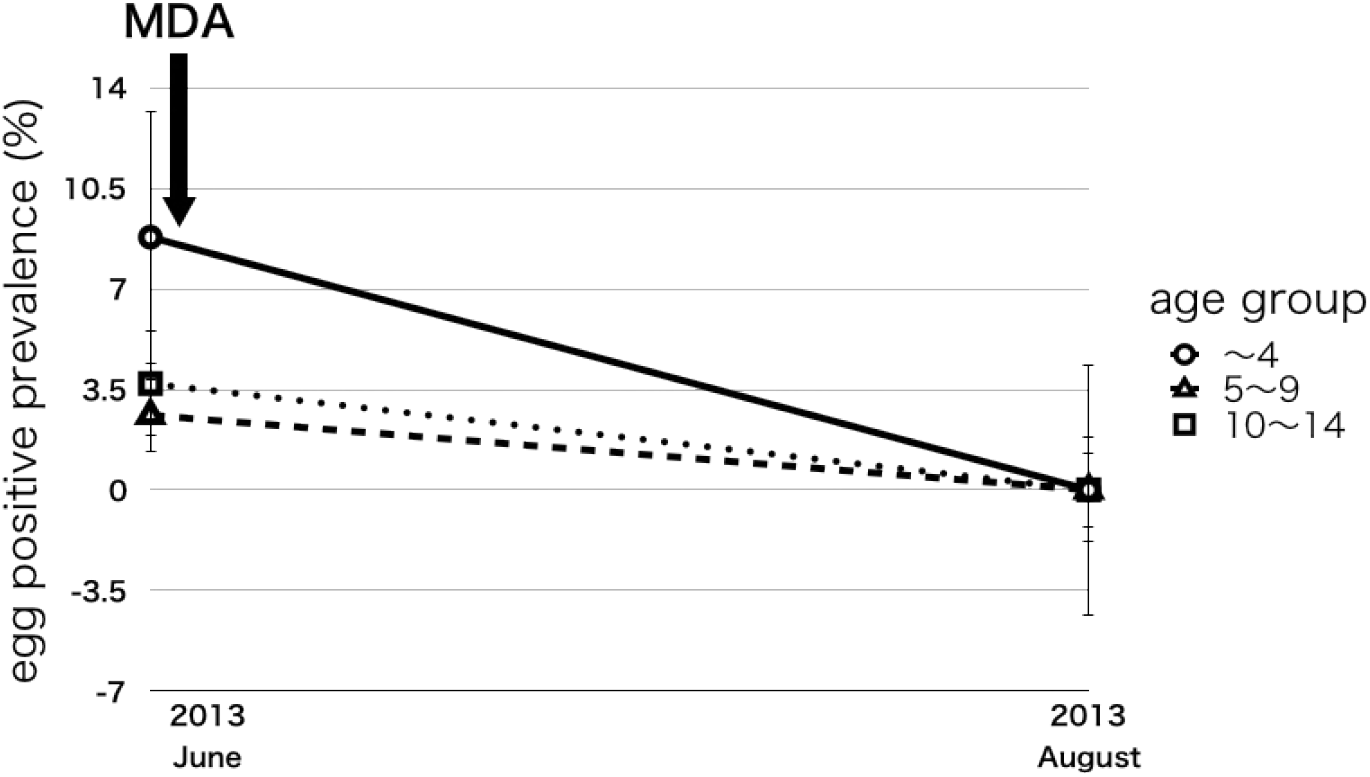
Process of egg positive prevalence by age group 2013

**Fig 4-b.**
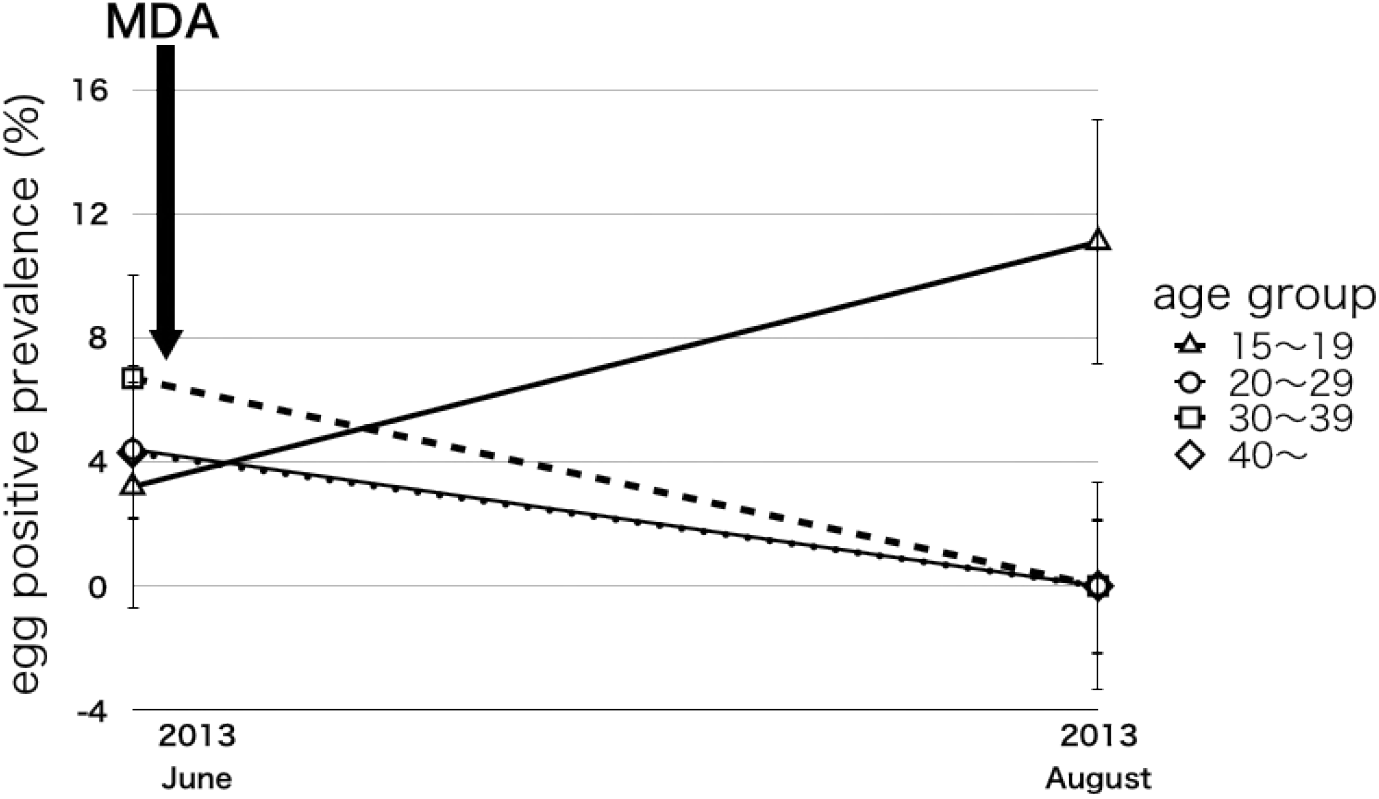
Process of egg positive prevalence by Age group 2013

**Table 2.** MDA Coverage Rate by Area in 2012

| Socio-demographic characteristics Area | Proportional MDA Attendance by Area 2012 |  |  |
| --- | --- | --- | --- |
|  | Registered | Treated | Percentage Coverage (%) |
| Chisindo | 390 | 196 | 50.3 |
| Mtika | 331 | 284 | 85.8 |
| Mapiri | 296 | 192 | 64.9 |
| Chisaka | 376 | 200 | 53.2 |
| Total | 1,393 | 772 | 55.4 |

**Table 3.** Results of urine filtering examination 2013

|  |  |  | 2013 |  |  |  |  |  | P value |
| --- | --- | --- | --- | --- | --- | --- | --- | --- | --- |
|  |  |  | June |  |  | August |  |  |  |
|  |  |  | Number of Subjects | Before MDA* |  | Number of Subjects | 8 weeks after MDA* |  |  |
|  |  |  |  | Egg positive number | Egg positive prevalence (95%CI) |  | Egg positive number | Egg positive prevalence (95%CI) |  |
| Total |  |  | 264 | 10 | 3.8 (2.1-6.9) | 211 | 2 | 0.9 (0.3-3.4) | P=0.050 |
| Male |  |  | 99 | 6 | 6.1 (2.8-12.7) | 86 | 1 | 1.2 (0.2-6.4) | P=0.082 |
| Female |  |  | 165 | 4 | 2.4 (0.9-6.1) | 125 | 1 | 0.8 (0.1-4.5) | P=0.293 |
| Age group | ~4 | Male | 14 | 1 | 7.1 (1.3-31.8) | 8 | 0 | 0.0 (0.0-32.9) | P=0.439 |
|  |  | Female | 20 | 2 | 10.0 (2.8-30.4) | 15 | 0 | 0.0 (0.0-20.7) | P=0.207 |
|  |  | Total | 34 | 3 | 8.8 (3.0-23.1) | 23 | 0 | 0.0 (0.0-14.6) | P=0.143 |
|  | 5~9 | Male | 20 | 1 | 5.0 (0.9-23.9) | 22 | 0 | 0.0 (0.0-15.1) | P=0.288 |
|  |  | Female | 19 | 0 | 0.0 (0.0-17.1) | 19 | 0 | 0.0 (0.0-17.1) | - |
|  |  | Total | 39 | 1 | 2.6 (0.4-13.3) | 41 | 0 | 0.0 (0.0-8.7) | P=0.302 |
|  | 10~14 | Male | 28 | 1 | 3.6 (0.6-17.9) | 22 | 0 | 0.0 (0.0-15.1) | P=0.371 |
|  |  | Female | 26 | 1 | 3.8 (0.7-19.1) | 33 | 0 | 0.0 (0.0-10.6) | P=0.256 |
|  |  | Total | 54 | 2 | 3.7 (1.0-12.7) | 55 | 0 | 0.0 (0.0-6.7) | P=0.150 |
|  | 15~19 | Male | 9 | 1 | 11.1 (2.0-43.9) | 8 | 1 | 12.5 (2.2-47.5) | P=0.929 |
|  |  | Female | 22 | 0 | 0.0 (0.0-15.1) | 10 | 1 | 10.0 (1.8-40.8) | P=0.132 |
|  |  | Total | 31 | 1 | 3.2 (0.6-16.4) | 18 | 2 | 11.1 (3.1-33.1) | P=0.267 |
|  | 20~29 | Male | 13 | 1 | 7.7 (1.4-33.6) | 15 | 0 | 0.0 (0.0-20.7) | P=0.274 |
|  |  | Female | 32 | 1 | 3.1 (0.5-15.9) | 20 | 0 | 0.0 (0.0-16.4) | P=0.425 |
|  |  | Total | 45 | 2 | 4.4 (1.2-15.0) | 35 | 0 | 0.0 (0.0-10.1) | P=0.207 |
|  | 30~39 | Male | 3 | 1 | 33.3 (6.1-79.5) | 2 | 0 | 0.0 (0.0-66.2) | P=0.361 |
|  |  | Female | 12 | 0 | 0.0 (0.0-24.6) | 5 | 0 | 0.0 (0.0-43.9) | - |
|  |  | Total | 15 | 1 | 6.7 (1.2-30.1) | 7 | 0 | 0.0 (0.0-35.9) | P=0.484 |
|  | 40~ | Male | 12 | 0 | 0.0 (0.0-24.6) | 9 | 0 | 0.0 (0.0-30.3) | - |
|  |  | Female | 34 | 2 | 5.9 (1.6-19.3) | 23 | 0 | 0.0 (0.0-14.6) | P=0.236 |
|  |  | Total | 46 | 2 | 4.3 (1.2-14.7) | 32 | 0 | 0.0 (0.0-10.9) | P=0.232 |

The cost of one urine reagent strip and one tablet of praziquantel were US$0.06 and US$0.125 in 2013 in Malawi. After MDA in reducing the egg positive prevalence, the positive predictive value was decreased and the negative predictive value was more than 96%. This suggests that it is practical to exclude urine occult blood negative subjects from receiving praziquantel.

## Discussion

Agriculture is one of main source of employment for people in developing countries such as Malawi. In fact, agriculture produces employment for more than 80% of the active labour force in Malawi [7]. The International Labour Organization (ILO) states that agricultural work is one of the most hazardous occupational activities to health worldwide [14]. Many kinds of agricultural activities are related to occupational injuries and those who are engaged in agriculture are being exposed to the risk of schistosome infection because they come into contact with freshwater during farm work in schistosome endemic countries. Although schistosomiasis is not specifically documented in the list of ILO occupational diseases [15], but it should be related to the occupational risks. Schistosomiasis is associated with populations living in poverty in sub-Saharan Africa, including Malawi. In order to ameliorate poverty, it is key to improve the working environment and maintain sustainable economic growth. It is because that good health significantly promotes economic growth, both in the short-run and long-run [16]. The labour force age ranges from 15-to 64-years of age, and 15-to 29-year olds are presumed to be the main labour force generation, accounting for 51.6% among all labour force generation [17]. Schistosomiasis could be presumed to have concerning with occupational risks in sub-Saharan African countries where agriculture is the main industry. If schistosomiasis has occupation-related risks, then adequate healthcare service should be provided for the labour force in order to provide a stable GDP growth rate. Improving health conditions boosts the productivity of workers and that increases economic growth in the long-run [16]. Thus, it is expected that protecting the health of the labour force could lead to a reduction in poverty in tropical and subtropical countries and mobilize national development in the region.

Schistosomiasis is prevalent throughout Malawi. Since the late 1990s, the decline in human capital accelerated the collapse of public health services [18]. The country depends on the income generated from agriculture. More than half of the Malawi population is food insecure [19] and 65.3% of the people are unable to meet their daily dietary needs [20]. According to previous studies [7–10, 21–24], school-aged children are a high-risk group for schistosomiasis infection. Our study showed that before the first MDA in June 2010, those in the twenties showed the highest egg positive prevalence (Table 1). Those in the twenties belonged to the labour force. To alleviate poverty, and cases of schistosomiasis related to poverty, it is important for the economic growth and development of this country to protect the health of the labour force. In the older-than-20 age group, the egg positive prevalence decreased ten months after the first MDA (Fig 2–b). In general, providing health information about schistosomiasis may bring about behavioral changes in the population that would improve the overall health in the country.

An increase in the egg positive prevalence was observed in the under-15-age group one year after MDA in June 2011. Among under-15-age group, the gradient of the increasing line of under-5 age-group was steepest (Fig 2a). Positive egg prevalence of each age group in 2013 was lower than the average in Malawi, and the highest was 8.8% in the under-5-age group (Table 3). There is a high risk of infection for those under the age of 15 years, and it is suggested that this tendency may be greater among those under the age of 5 years. Previous studies reported that pre-school children are also at the risk of schistosome infection [25], and when school-aged children were screened schistosome infection ranged from 5% to 57% [26]. In order to confirm the minimum age of schistosome infection, all subjects, of all ages, in the study underwent a urine examination in 2013. We detected that the age of the youngest infected subject was 2-years-old in this study. Pre-school children frequently accompany their guardians into the freshwater areas [27]. Although it is known that pre-school children also face schistosome infection [28], there is still room for further study on the safety of administering praziquantel to children less than 4 years of age. MDA, however, may be a more promising approach to disease control in Malawi [29]. The Malawi National Schistosomiasis Control Programme does not have well-documented evidence of the universal drug treatment [30]. As the first step in breaking the chain of chronic and repetitive infection, the first year of school enrollment is considered an appropriate time for the first MDA after birth.

In only 15-19 age group, elevation of the egg positive prevalence was confirmed after MDA in August 2013 (Fig 4b). Therefore, those who graduated from schools –who are in the 15-19 age group-belong to the high-risk group for repeat schistosome repeated.

Referring to the population pyramid of Malawi in 2010, teens, twenties, and school-aged children accounted for about 60% of the total population [31]. The population of teens and twenties is more than four million. These two generational groups are presumed to be major labour force; so if their health suffers due to schistosome infection, it may have undesirable effects on national development and economic growth. As previously mentioned, agriculture is the primary industry and provides more than 80% of employment in the country [7]. Moreover, community-wide MDA of praziquantel is highly cost effective when compared with treatment of school-aged children alone [32]. Therefore, it is important to protect the health of not only school-aged children, but also the overall labour force. There seems to be a link between health and income growth in the schistosomiasis endemic areas. The main route of schistosome infection is through contact with infected freshwater. The daily activities of Malawians result in contact with freshwater through fishing, farming, washing, bathing, swimming, and so on. The occupations which are at risk of infection with schistosomiasis in Malawi are; rice farmers, sugarcane growers, irrigators thus those responsible for opening water flow in canals, fishermen, tobacco growers, vegetable farmers, cattle wrestlers, held man, cane cutters, fish pond workers, and wildlife guards [9]. For instance in Japan, a former schistosomiasis endemic country (*Schistosoma japonicum*), schistosomiasis was regarded as an occupational disease for rice farmers [33]. And farmers’ health injuries caused by schistosomiasis were symbolically elaborated in Katayama Memoir (Katayama-ki) written by Dr Yoshinao Fujii in 1847. Human excrement, often containing schistosome eggs, is spread in fertilizing the fields, and the barelegged farmers in rice-paddies are easy victims for cercariae. Generally, urination and defecation are the main methods for schistosome eggs to get into the environment. The transmission of schistosomiasis in Malawi remains fragmented [34], and setting up proper toilets and designating specific places for latrines/toilets in each district is thought to be an effective method of infection control. From 2010 to 2011 survey, the egg positive prevalence after MDA decreased, but it was not eradicated. On the other hand, in the survey of 2013, egg positive prevalence was reduced to 0% in all age groups except in the 15-19 age group (Figs 4-a and b). The surveyed areas in 2013 are located near the Capital City Lilongwe, where health education and promotion about schistosomiasis infection was conducted at schools and there were frequent media broadcasts on the subject. These activities may have led to differences in the survey results. Workers in rural areas are more likely to earn less than their counterparts in urban areas [15]. The difference in income may also have a bearing on schistosome infection. In Nkhotakota district on Lake Malawi, it is assumed that many residents are in contact with fresh water area more frequently because agricultural activities are greater than in the environs of Lilongwe.

Haematobium schistosomiasis can cause and aggravate anemia caused by low dietary iron, hookworm infection, and malaria in Malawi. The health hazards related to haematobium schistosomiasis can include bladder cancer and cervical cancer, but most likely there are repeated infections and chronic infections occurring with these severe diseases. Although bladder cancer occurs principally as urothelial carcinoma, the major histological cell type of bladder cancer associated to haematobium schistosomiasis is squamous cell carcinoma [35–37]. Because the occurrence of squamous cell carcinoma is associated with persistent chronic inflammation, it is essential to prevent chronic infections and repetitive infections to suppress its development.

MDA of praziquantel was conducted in June 2010, and the egg positive prevalence one year later decreased to about two fifth as shown in Fig 1 (p<0.01). Egg positive prevalence in the target areas in 2013 where MDA was conducted in 2012 was 3.8%; and it was supposed to be lower than previously reported egg positive prevalence in Malawi. These results showed that MDA of praziquantel could certainly reduce egg positive prevalence of haematobium schistosomiasis. Although MDA is not an effective replacement for the existing vector control, MDA has the potential to reduce transmission for a limited time and has to be repeated regularly for sustained effect [38]. However, the egg positive prevalence in August 2013 was reduced from 3.8% to 0.9% two months after MDA (p=0.050). This result indicates that the effectiveness of annual MDA might reduce over time. Thus, it may not be necessary to carry out MDA every year. After mass drug administration of praziquantel, no resident showed serious adverse reactions and is also considered safe and appropriate.

Former studies indicated the cost-effectiveness of urine reagent strips [39, 40], but it is still uncertain whether the urine reagent strip is cost-effective [41, 42]. We analyzed the effectiveness of the urine reagent strips for checking hematuria [43]. The cost of one reagent strip and one tablet of praziquantel was US$0.06 and US$0.125 in 2013 in Malawi. After MDA in reducing the egg positive prevalence, the positive predictive value was decreased and the negative predictive value was more than 96%. This suggests that it is practical to exclude urine occult blood negative subjects from receiving praziquantel; and then this screening for MDA could lead to cost-effectiveness for schistosomiasis control. Since the urine reagent strips can also produce false negatives, continuation of MDA is important for achieving better infection control.

Assuming 1,810 subjects for MDA of praziquantel and presupposing the average body weight of the subject to be 40kg, three tablets of praziquantel are needed to for each subject. Under this assumption, if praziquantel is administered to all 1,810 subjects, the total cost will be US$678.80, US$0.38 per person. However, assuming an egg positive prevalence of 40% and screening using urine reagent strips and administering praziquantel, the total cost can be estimated to be US$515.90, US$0.29 per person. If the egg positive prevalence is 40%, screening subjects for MDA using urine reagent strips, the cost reduction can be estimated to be about 24% -showing an overall cost reduction. It should be noted that in a very-low egg positive prevalence settings, microhaematuria is an unstable manifestation for haematobium schistosomiasis and the treatment decision should not be based on the urine reagent strips results alone [44]; different kinds of examinations for differential diagnoses should be performed according to each disease condition. Despite the reported high rate of infection noted in the previous study, the tendency to seek medication from a medical facility is not substantial, with only 34.7% of the respondents seeking treatment for haematuria at the nearest medical facility [45]. If someone has symptoms but leaves the facility, the chain of chronic/repetitive schistosome infection can not be broken. Therefore, when symptoms such as hematuria are presented, it is important to thoroughly inform every resident that he/she should consult a medical institution promptly without neglecting it. Sustainable health education for children and young adults is the pillar for controlling schistosome infection and it could lead to better health management.

Schistosomiasis, without proper intervention and treatment, belongs to a group of diseases that could lead to morbidity and mortality to the residents in schistosome endemic countries. And yet, schistosomiasis does not draw as much attention as malaria and tuberculosis, it is still one of the neglected tropical diseases (NTDs). Not enough countermeasures are taken to control the disease; and it could choke off economic development and growth in low-income tropical countries such as Malawi. Although it is expected that urgent schistosomiasis countermeasures will make a great social contribution in affected tropical countries, it can be easily imagined that the budgeting for countermeasures will be quite difficult in those countries. Thus, it is considered essential to prioritize to the implementation of countermeasures. School-aged children were the main targets for schistosomiasis countermeasure in the past, but this research shows that both the infection rate and the recurrence rate were higher in the labour force and that may have a direct influence on economic development. Agricultural work is the main form of labour in many developing countries, including Malawi, and contact with freshwater areas is inevitable as long as the residents are engaged in agricultural activities. Therefore, schistosomiasis should be considered to have concerning with occupational risks. Good health is positively related to economic growth or output [16]. Based on the results of this study, we believe that reasonable countermeasures and well-targeted treatment could reduce the prevalence of haematobium schistosomiasis; and this could lead to an improvement in morbidity and mortality, reducing the prevalence of schistosomiasis in Malawi and other schistosome endemic countries. It will be resulted to protect health of labour force, too. Furthermore, since schistosomiasis is presumed to have occupation-related risks, we consider that schistosome control will be a valuable step-up to economic development and make a social contribution in Malawi and many low-income tropical countries.

## Acknowledgement

This research was supported by <u>JSPS KAKENHI Grant Number JP23406025</u>. And Ms. Yuko Harada (Nippon International Cooperation for Community Development, Japan: NICCO) provided eager support and made a great contribution to our research.

